# Self-confocal NIR-II fluorescence microscopy for in vivo imaging

**DOI:** 10.1101/2021.12.19.473402

**Authors:** Jing Zhou, Tianxiang Wu, Liang Zhu, Yifei Li, Liying Chen, Jun Qian

## Abstract

Benefiting from low scatter of NIR-II light in biological tissues and high spatial resolution of confocal microscopy, NIR-II fluorescence confocal microscopy has been developed recently and achieve deep imaging in vivo. However, independence of excitation point and detection point makes this system difficult to be adjusted. New, improved, self-confocal NIR-II fluorescence confocal systems are created in this work. Based on a shared pinhole for excitation light and fluorescence, the system is easy and controlled to be adjusted. The fiber-pinhole confocal system is constructed for cerebrovascular and hepatocellular NIR-II fluorescence intensity imaging. The air-pinhole confocal system is constructed for cerebrovascular NIR-II fluorescence intensity imaging, hepatic NIR-II fluorescence lifetime imaging, and hepatic multiphoton imaging.

## 1. Introduction

Traditional visible-light fluorescence confocal microscopy is an efficient method for high-resolution imaging of cells ^[1]^. However, visible light has large scatter in biological tissues, leading it can not penetrate deeply, and thus the imaging depth is very limit ^[2]^. Light in the near infrared two spectral region (NIR-II, 900-1700 nm) with less scatter can over come this problem in a certain extent ^[3, 4]^. Combining it to the confocal system, can achieve deep imaging depth with high spatial resolution in vivo ^[5, 6]^. Although NIR-II fluorescence confocal system has been developed, it has some annoying point. Adjusting the pinhole at the detection modular to conjugate with the excited point in the sample is hard and very time-consuming. Since the NIR-II fluorescence signal is weak and invisible, so the position of the pinhole is unpredictable and the adjustment process is difficult to control. Another important point, the quantum efficiency of NIR-II detector, normally InGaAs PMT, is very low. So the requirement for conjugation of excitation and detection points is very strict.

In our work, new, improved, NIR-II confocal systems, are proposed. One physical pinhole is simultaneously as excitation point and detection point, as long as the excitation light emits from this pinhole-point, the fluorescence must will come back to and focused on this pinhole-point again. Depending on its self-confocal characteristic, the system becomes easily and controllably to be adjusted and maintained. A non-adjustable fiber-pinhole system adapted to continuous wave (CW) laser was constructed for NIR-II fluorescence intensity imaging. A conveniently-adjustable air-pinhole system adapted to femtosecond (fs) pulsed laser was constructed for NIR-II fluorescence intensity imaging, NIR-II fluorescence lifetime imaging and multiphoton imaging.

## 2. Results

### 2.1 Fiber-pinhole self-confocal NIR-II fluorescence microscopy

The fiber-pinhole self-confocal NIR-II fluorescence microscope using a fiber core as the pinhole, so it is adapted to CW laser excitation, obtaining the intensity information of fluorophore (Fig. 1). The optical fiber and the collimator in the dashed box of Fig.1 compose the self-confocal module. The excitation light (800 nm CW laser) passing through the short-pass dichroic mirror (DMSP) be focused on the optical fiber core by a collimator, and then propagate to another optical fiber core (fiber-pinhole, diameter = 105 um), expanded by a collimator and incident on the mirror. After exciting the fluorophore, the fluorescence will roughly come back along the excitation path, and refocus on the fiber-pinhole again. Then the fluorescence will be reflected by the dichroic mirror and be collected by the photomultiplier tube (PMT). This process is automatic and non-adjustable, saving time and improving efficiency to a great extent.

**Fig. 1.**
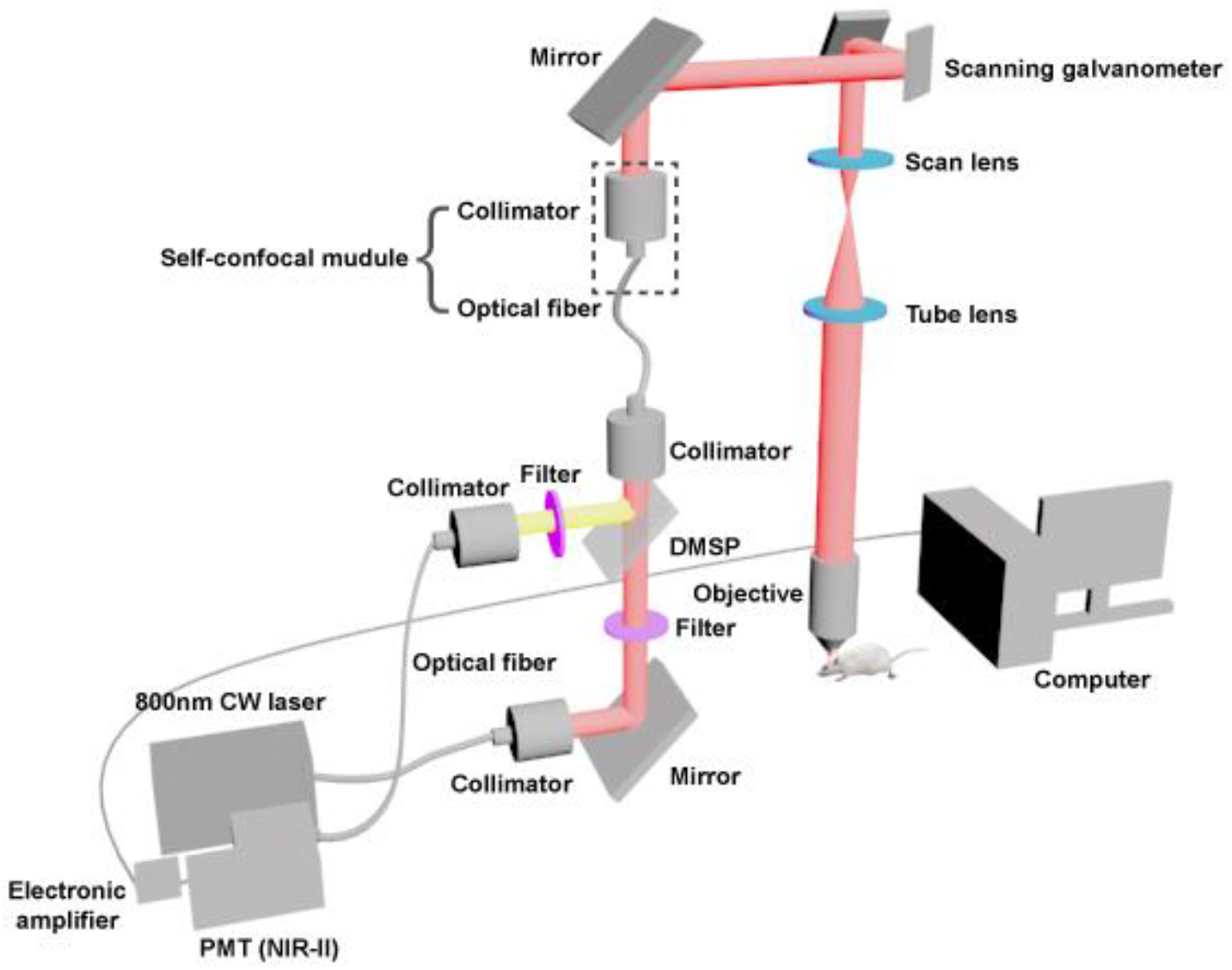
Schematic of fiber-pinhole self-confocal NIR-II fluorescence microscope.

We used this system to image cerebral vessels of mice intravenously injected with indocyanine green (ICG) in vivo. ICG’s absorption peak is around 800 nm. And its fluorescence tail is beyond 950 nm (Fig. S1). The imaging depth is ∼ 700 μm (Fig. 2). Diameters of vessels marked by dotted lines at 200 μm depth (Fig. 2c), 350 μm depth (Fig. 2e) and 600 μm depth (Fig. 2h) are 4.7 μm, 3.4 μm and 3 μm respectively. The image resolution is high.

**Fig. 2.**
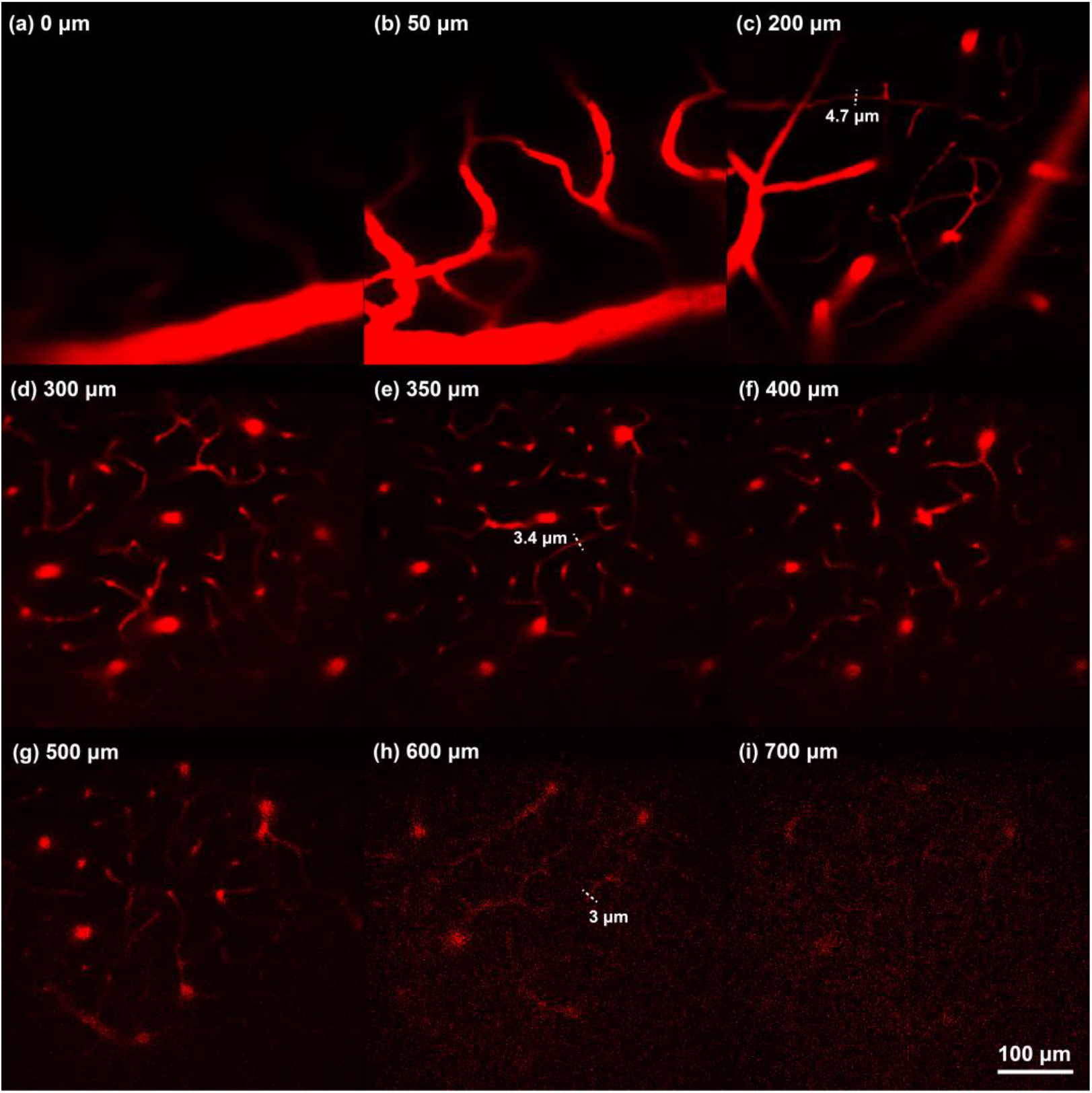
Fiber-pinhole system for cerebrovascular NIR-II fluorescence intensity imaging. (a-i) Cerebrovascular images at various depths from 0 μm to 700 μm. The mouse was injected with ICG (5 mg/mL, 200 μL). Excitation: 800 nm CW laser (60 mW before objective); fluorescence collection range: > 950 nm; scanning speed: 10 us/pixel, PMT voltage: 500 V. Scale bar: 100 μm.

We also used this system to image hepatocytes of mice intravenously injected with ICG in vivo, since ICG is metabolized by digestive system, so it will be ingested by hepatocytes ^[7]^. Hepatocyte can be clearly imaged by this system with small depth of focus (Fig. 3a). Cytoplasm and nucleus are distinct. Along the dotted line on the hepatocyte, the intensity distribution is shown in Fig. 3b.

**Fig. 3.**
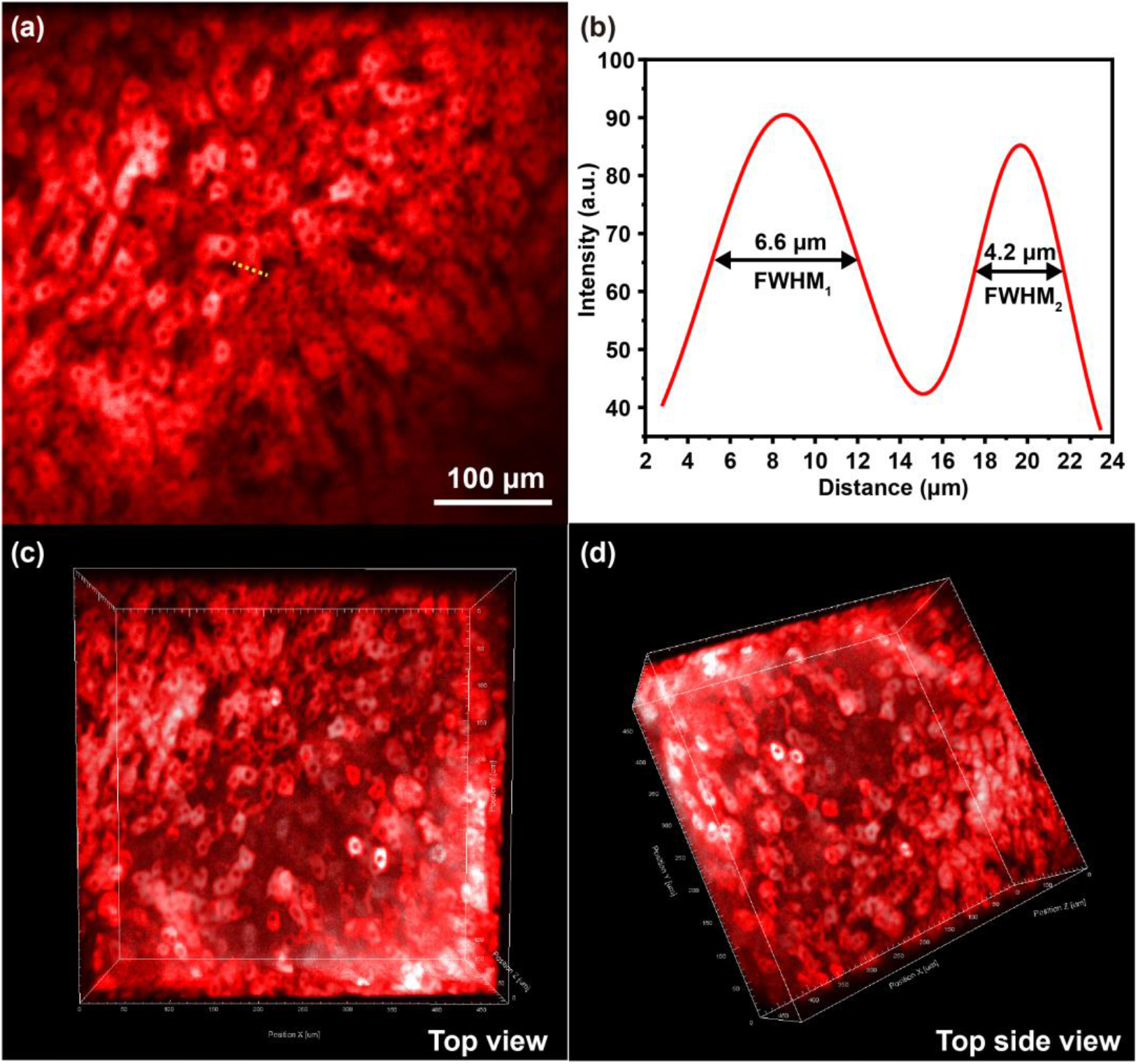
Fiber-pinhole system for hepatocellular NIR-II fluorescence intensity imaging. (a) NIR-II confocal imaging of hepatocytes of a mouse. The mouse was injected with ICG (5 mg/mL, 200 μL). Excitation: 800 nm CW laser (5 mW before objective); fluorescence collection range: > 950 nm; scanning speed: 10 us/pixel, PMT voltage: 450 V. Scale bar: 100 μm. (b) The intensity distribution along the dotted line in (a) on a hepatocyte taking in ICG and the corresponding FWHM analyses. (c, d) Top view and top side view of hepatocellular 3D reconstruction.

The cell’s width is ∼ 20 μm. FWHMs of peaks represent widths of cytoplasm, 6.6 μm and 4.2 μm. The total imaging depth is 184 μm, and the 3D reconstructions of superficial liver at top view and top side view are shown (Fig. 3c and 3d).

### 2.2 Air-pinhole self-confocal NIR-II fluorescence microscopy

The air-pinhole self-confocal NIR-II fluorescence microscope using a diaphragm with continuously adjustable aperture as the pinhole, so it can be adapted to fs pulsed laser excitation, obtaining fluorescence intensity information and lifetime information of fluorophore, and can achieve simultaneously NIR-II fluorescence confocal imaging and multiphoton imaging (Fig.4). Lens_ex_, pinhole and lens_fl_ in the dashed box of Fig.4 compose the self-confocal module. The excitation light (800 nm fs laser) passing through the short-pass dichroic mirror be focused on the air-pinhole by lens_ex_, and then be expanded by lens_fl_ and incident on the mirror. After exciting the fluorophore, the NIR-II fluorescence will roughly come back along the excitation path, and refocus on the air-pinhole again. Then the NIR-II fluorescence will be reflected by the dichroic mirror and be collected by the PMT (NIR-II). With pulsed excitation light and time-correlated single-photon counting (TCSPC) module in the computer, the NIR-II fluorescence lifetime can be calculated. In addition, 800 nm fs laser can excite the multiphoton fluorescence of the fluorophore, which will be reflected by the long-pass dichroic mirror (DMLP) and collected by the PMT response to the visible light. So NIR-II fluorescence confocal imaging and multiphoton imaging can be carried simultaneously for multi-structure observation. This system is conveniently-adjustable and multifunctional, saving time and improving efficiency to a large extent.

**Fig. 4.**
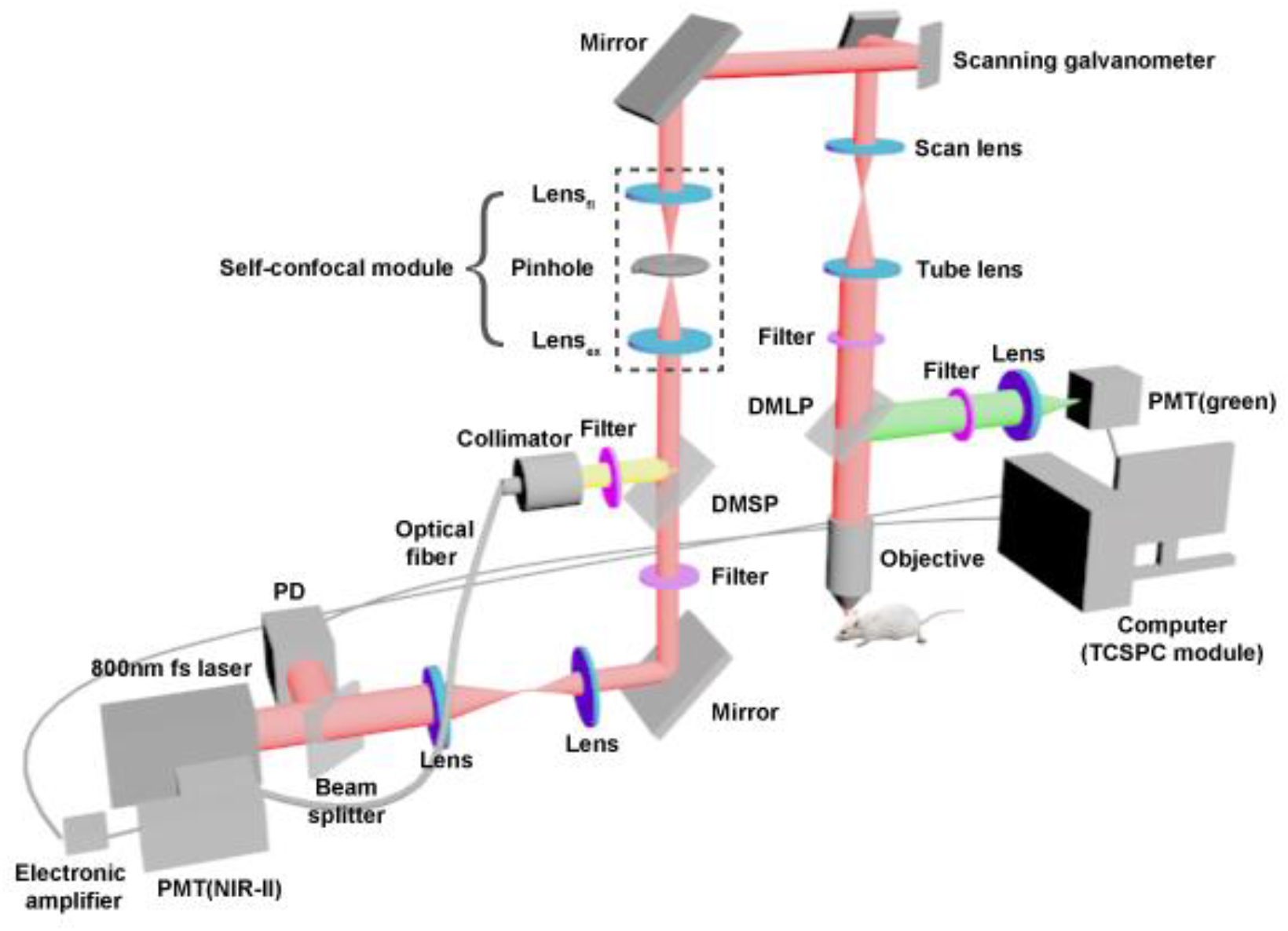
Schematic of air-pinhole self-confocal NIR-II fluorescence microscope.

We used this system to do intensity imaging of cerebral vessels of mice intravenously injected with ICG in vivo. The imaging depth is ∼ 760 μm (Fig. 5). Diameters of vessels marked by dotted lines at 200 μm depth (Fig. 5c), 600 μm depth (Fig. 5g) and 700 μm depth (Fig. 5h) are 3.6 μm, 5.4 μm and 4.8 μm respectively. The image resolution is high. The 3D reconstructions at top view and top side view are shown (Fig. 5j and 5k).

**Fig. 5.**
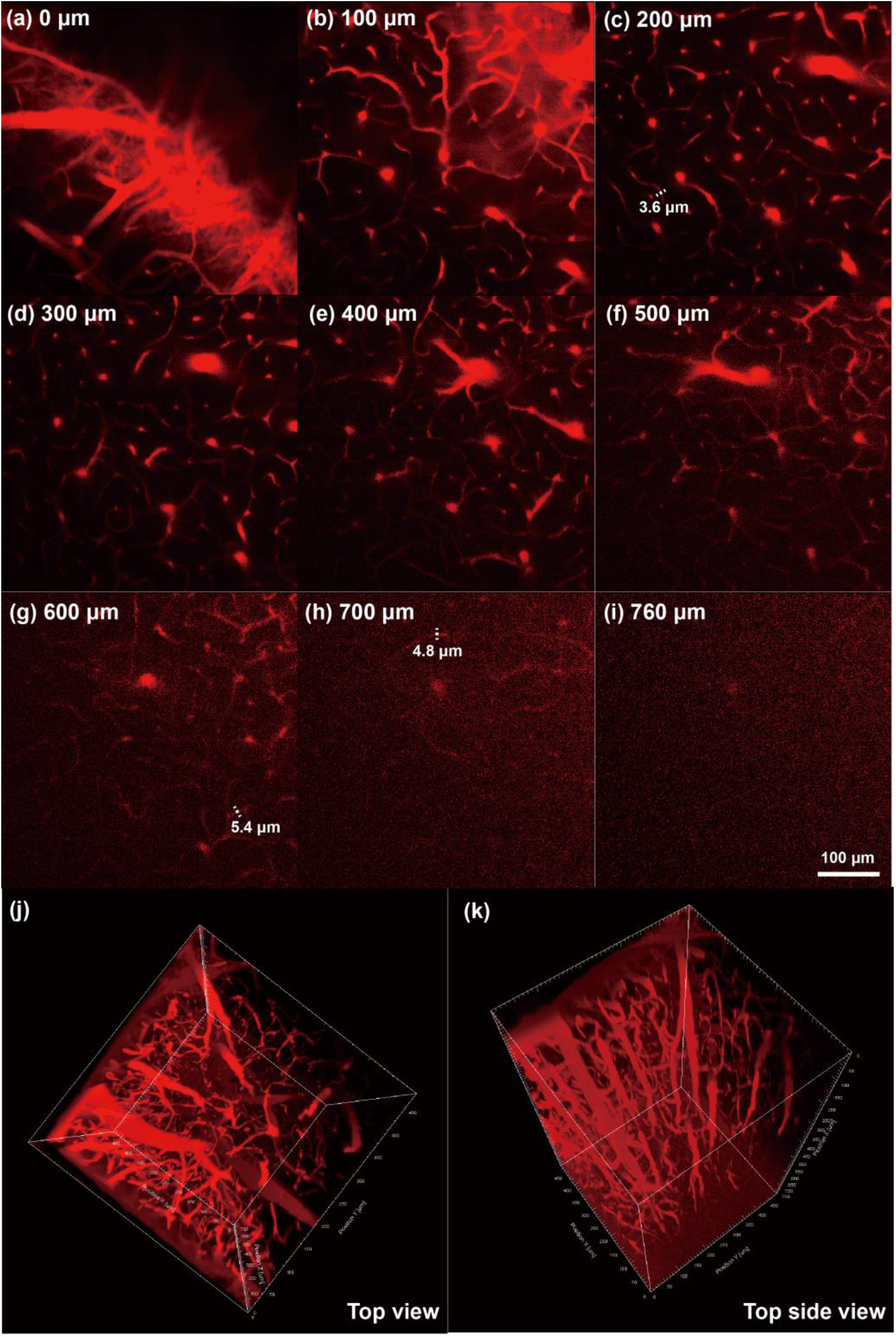
Air-pinhole system for cerebrovascular NIR-II fluorescence intensity imaging. (a-i) cerebrovascular images at various depths from 0 μm to 760 μm. The mouse was injected with ICG (5 mg/mL, 200 μL). Excitation: 800 nm fs pulsed laser (80 mW before objective); fluorescence collection range: > 950 nm; pinhole aperture: 400 μm; scanning speed: 10 us/pixel, PMT voltage: 600 V. Scale bar: 100 μm. (j, k) Top view and top side view of cerebrovascular 3D reconstruction.

Utilizing fluorescence lifetime measurement function of this system, we measured fluorescence lifetime of two fluorophores in vitro, ICG and 2TT-oC26B NPs ^[9]^, at first. ICG’s lifetime is ∼ 689 ps, while 2TT-oC26B NPs’ is ∼ 404 ps (Fig. 6a and 6b). After injecting the ICG into the mouse, it will be distributed in hepatocytes. While injecting 2TT-oC26B NPs to the mouse, it won’t be in hepatocytes, but in other hepatic components. We imaged livers of two mice intravenously injected with ICG and 2TT-oC26B NPs respectively. The lifetime of ICG in hepatocytes (blue) is ∼ 700 ps, while that of 2TT-oC26B NPs in other liver components (orange) is ∼ 400 ps (Fig. 6c and 6d). They are coincident to the results in vitro. When injecting ICG and 2TT-oC26B NPs together to a mouse, with only one collected spectral range (> 950 nm) and one PMT, hepatocytes taking in ICG (blue) and other liver components marked by 2TT-oC26B NPs (orange) can still be distinctively distinguished by different fluorescence lifetime. The intensity images can be seen in Fig. S2. So this system is cost-effective and convenient for multi-structure imaging simultaneously.

**Fig. 6.**
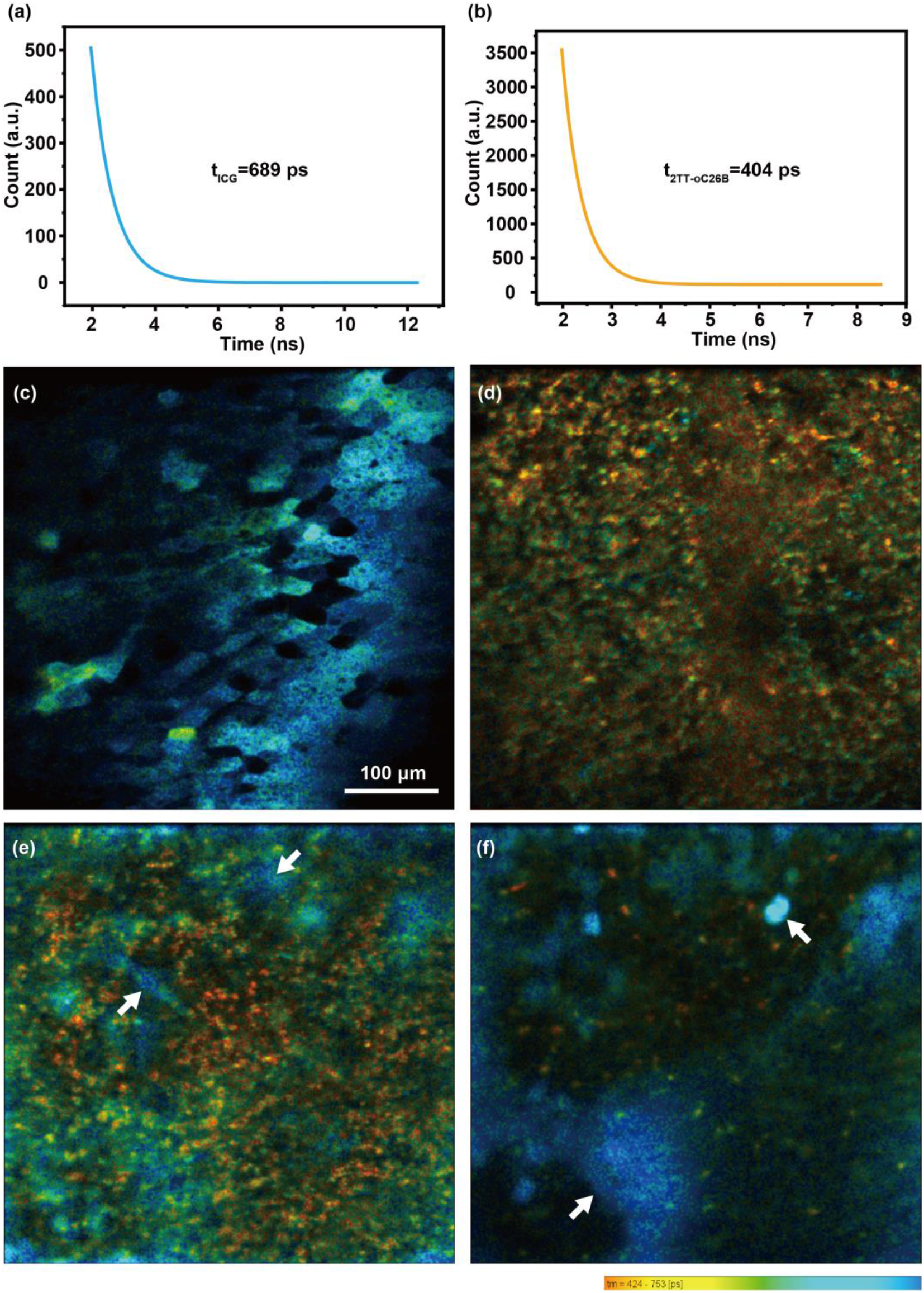
Air-pinhole system for hepatic structures NIR-II fluorescence multi-lifetime imaging. (a) Fluorescence lifetime of ICG in vitro. (b) Fluorescence lifetime of 2TT-oC26B NPs in vitro. (c) Fluorescence lifetime imaging of hepatocytes of a mouse intravenously injected with ICG (1mg/ml, 100 μL). (d) Fluorescence lifetime imaging of hepatic components of a mouse intravenously injected with 2TT-oC26B NPs (2mg/ml, 100 μL). (e, f) Fluorescence lifetime imaging of hepatocytes (blue) and other hepatic components (orange) of a mouse intravenously injected with ICG (1mg/ml, 100 μL) and 2TT-oC26B NPs (2mg/ml, 100 μL) at different depth. White arrows point to hepatocytes. Excitation: 800 nm fs pulsed laser (5 mW before objective); fluorescence collection range: > 950 nm; pinhole aperture: 400 μm; scanning speed: 10 us/pixel, PMT voltage: 500 V. Scale bar: 100 μm.

Benefiting from the air-pinhole, the fs pulsed laser won’t be broadened, and thus can excite multiphoton fluorescence. The liver of mouse has multiphoton autofluorescence (green light) under 800 nm fs laser excitation (Fig. 7a and 7d). Injecting ICG to the mouse, we can see NIR-II fluorescence confocal imaging of blood vessels in liver and hepatocytes (Fig. 7b and 7e). Meanwhile, in green spectral region, multiphoton autofluorescence of specific hepatic component can be detected. Fused images indicate that the specific hepatic component locates in the gap between hepatocytes, which may be blood, bile, or immune cells (Fig. 7c and 7f). Relying on this system, microscopic multi-channel imaging can bring into full play with conveniently-adjustable characteristic.

**Fig. 7.**
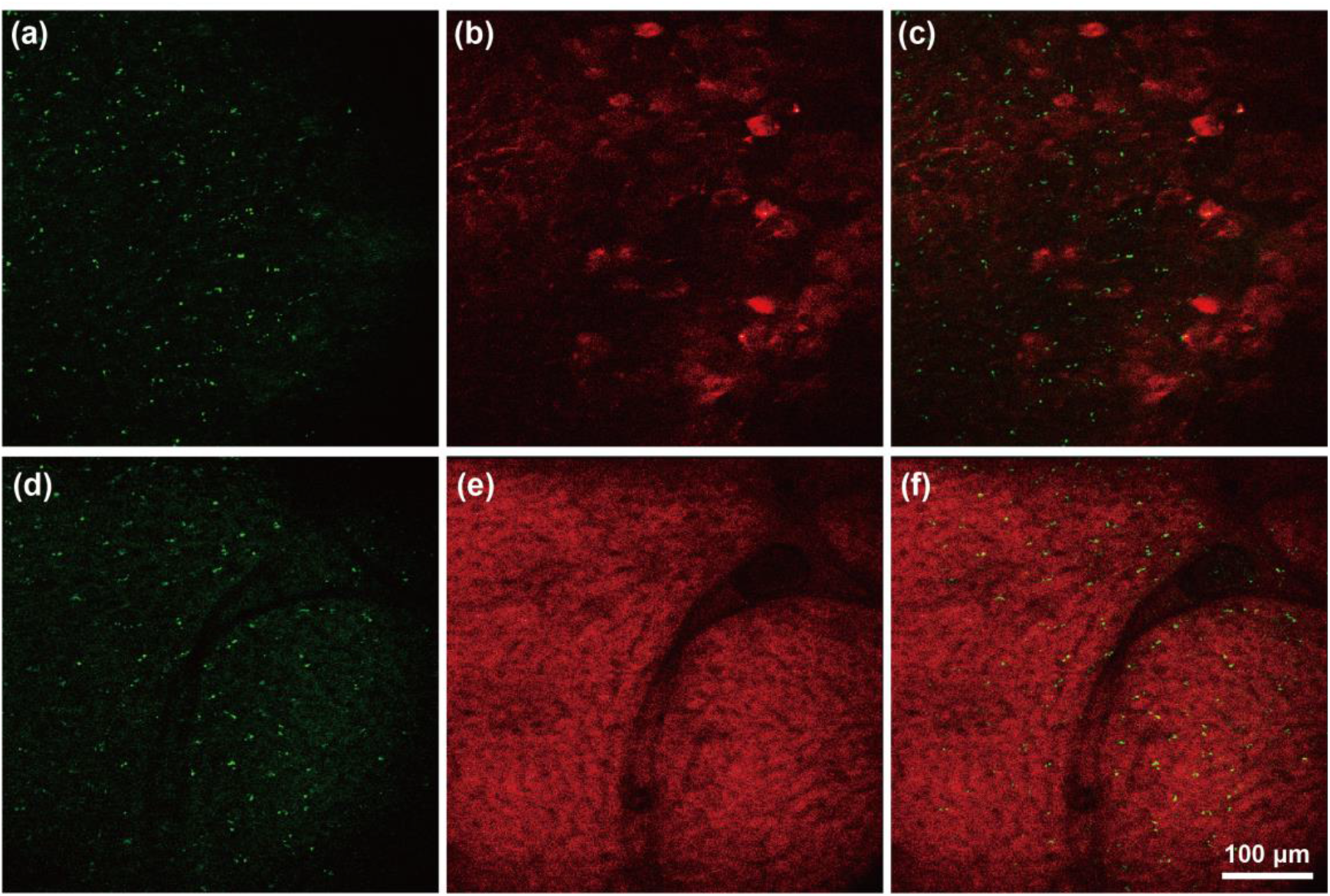
Air-pinhole system for hepatic structures NIR-II and multiphoton multi-mode imaging. (a, d) Multiphoton images of liver of the mouse in different fields. (b) NIR-II fluorescence confocal image of blood vessels in liver and hepatocytes of the mouse injected with ICG (1 mg/ml, 100 μL). (c) Fused image of (a) and (b). (e) NIR-II fluorescence confocal image of hepatocyte cord of the mouse injected with ICG (1 mg/ml, 100 μL). (f) Fused image of (d) and (e). Excitation: 800 nm fs pulsed laser (3 mW for NIR-II confocal imaging, 45 mW for multiphoton imaging before objective); fluorescence collection range: > 950 nm; pinhole aperture: 400 μm; scanning speed: 10 us/pixel, PMT voltage: 600 V. Scale bar: 100 μm.

## 3. Discussion

The self-confocal NIR-II fluorescence microscopy with the advantages of non-adjustable or conveniently-adjustable and deep imaging depth is charming. Fiber-pinhole self-confocal NIR-II fluorescence system adapted to CW laser can be used in normal intensity imaging, such as cerebrovascular imaging, hepatocyte imaging, et al, for observing structures. Air-pinhole self-confocal NIR-II fluorescence system adapted to fs laser can be used in not only intensity imaging, but also lifetime imaging and multiphoton imaging. Different fluorophores with different fluorescent lifetimes targeting different goals can be distinguished by the lifetime, such as ICG and 2TT-oC26B targeting hepatocyte and other hepatic components respectively. Besides, fluorescence lifetime of fluorophore can reflect its concentration, polarity and viscosity affected by the environment. So that we can know the metabolism of different fluorophores for evaluating the biosafety and pharmacokinetics, and know the cell cycle ^[8]^. Label-free or fluorophore-labeled multiphoton imaging is an advanced method for high-resolution and deep-depth imaging. Combining it with confocal NIR-II fluorescence microscopy can provide multi-channel imaging for observing multi-structure simultaneously, such as hepatocyte and other hepatic components, neurons with GFP and blood vessels, et al. What’s more, intensity imaging, lifetime imaging and multiphoton imaging can be integrated together to implement multi-plus-channel and multi-function imaging. These systems will effectively provide platform for biomedical researches, significantly promoting the biomedical field.

## 4. Materials and Methods

### 4.1 Materials

ICG was purchased from DanDong Pharmaceutical Factory (Liaoning, China). 2TT-oC26B NPs were synthesized according to our previous report ^[9]^. Phosphate buffered saline (PBS) was purchased from Sinopharm Chemical Reagent Co., Ltd., China.

### 4.2 Animal preparation

C57BL/6 mice (female, 6 weeks old) were used for in vivo experiments. They were provided by the Zhejiang Academy of Medical Sciences and kept at the Experimental Animal Center of Zhejiang University. The room temperature of the rearing environment was maintained at 24 °C with a 12 h light/dark cycle. Mice were continuously supplied with water and standard laboratory chows.

In the cerebrovascular imaging experiments, the skull of the anesthetized mouse was opened by microsurgery. Then a thin round cover glass with double wings was attached to the mouse brain by dental cement. The purpose of doing this is to protect and flatten the brain of the mouse to ensure the quality of microscopic imaging. Before imaging, both wings were fixed on the mouse rack to immobilize mouse’s head, and the mouse was then intravenously injected with ICG.

In hepatic imaging experiments, the mouse was anesthetized at first, then shaving and laparotomy were performed to completely expose the liver. Next, glass slide bridge was put on the abdominal cavity with medical adhesive tape coating to cover sharp edges and increase friction. Then hold large liver lobe using two wet cotton swabs and place it on the slide. A cover slip (24 mm x 32 mm) was put on the liver lobe fixed by medical tapes to ensure that it is oriented as horizontal as possible for suiting water immersion objective. The cover slip should be in contact with the tissue without squeezing it ^[10]^. ICG was intravenously injected to the mouse before imaging, and 2TT-oC26B NPs were intravenously injected to the mouse 1 day before imaging.

### 4.3 Ethical approval

All in vivo experiments are approved by the Institutional Ethical Committees of Animal Experimentation of Zhejiang University (ZJU20210254), and strictly abided by “The National Regulation of China for Care and Use of Laboratory Animals”.

### 4.4 Experimental set up for measuring absorption spectrum and fluorescence spectrum

The absorption spectrum of ICG was measured by UV-VIS-NIR spectrophotometer (CARY 5000, Agilent). The fluorescence spectrum of ICG was acquired on a home-built system based on the PG2000 spectrometer (370 nm - 1050 nm, Ideaoptics Instruments) and NIR2200 spectrometer (900 nm - 2200 nm, Ideaoptics Instruments).

### 4.5 Experimental set up of fiber-pinhole self-confocal NIR-II fluorescence microscopy

800 nm CW laser (Maitsse, continuous single longitudinal mode tunable; average output power @ 800 nm: 3 W; Spectra-Physics) beam is coupled into an optical fiber and collimated by a collimator. Then the laser incidents on a mirror for tuning the propagation direction of it and focused into the fiber-pinhole. After reflected by the mirror, laser passes through an 850 nm short-pass filter (FESH0850, Thorlabs) and a 950 nm short-pass dichroic mirror (DMSP950R, Thorlabs), and then be focused into the optical fiber by a collimator and propagates in the fiber to the fiber-pinhole (optical fiber core, diameter = 105 μm), then collimated by another collimator again. The collimated laser is guided into a commercial scanning microscope (FV1200, Olympus), incident to a mirror, then reflected to the scanning galvanometer (scanning speed: 10 μs/pixel; scanning area: 512 pixels × 512 pixels), which controls the scanning process of laser focal point in the x-y directions. After passing through the scan lens, tube lens and an objective (XLPLN25XWMP2, 25×, NA = 1.05, Olympus) with near infrared anti-reflection film, laser is eventually focused on the sample. NIR-II fluorescence from the sample passes back roughly along the excitation path, and refocus into the fiber-pinhole again. Propagating in the fiber and collimated by the collimator, the fluorescence is reflected by the 950 nm short-pass dichroic mirror (DMSP950R, Thorlabs) and passes through a 950 nm long-pass filter (FELH0950, Thorlabs). The fluorescence above 950 nm is then focused into an optical fiber by a collimator. Then, fluorescence signal guided in the fiber is focused onto a NIR (950–1700 nm) sensitive PMT (H12397-75, Hamamatsu). The electrical signal generated in the PMT is amplified by an electrical signal amplifier (C12419, Hamamatsu), based on which an image could be reconstructed via a computer.

### 4.6 Experimental set up of air-pinhole self-confocal NIR-II fluorescence microscopy

800 nm fs pulsed laser (Mira-HP; repetition rate: 76 MHz; Coherent) beam is divided into two beams by a beam splitter at first, one weak beam reflected by the beam splitter is detected by a photodiode (PD) to synchronize the time information of laser pulse. Another beam’s width is shrunk by a pair of lenses after passing through the beam splitter, and then is reflected by a mirror, passing through an 850 nm short-pass filter (FESH0850, Thorlabs) and a 950 nm short-pass dichroic mirror (DMSP950R, Thorlabs), then focused into the aperture of a diaphragm (air-pinhole, aperture-adjustable) by the lens_ex_. The excitation light point is then expanded by lens_fl_ and guided into the commercial scanning microscope (FV1200, Olympus). After expanded by the scan lens and tube lens, the excitation light passes through a 650 nm long-pass dichroic mirror (DMLP650R, Thorlabs) and is focused on the sample. NIR-II fluorescence from the sample passes back roughly along the excitation path, and refocus into the air-pinhole again. Collimated by the lens_ex_ and reflected by the 950 nm short-pass dichroic mirror (DMSP950R, Thorlabs), NIR-II fluorescence signal is finally detected by the NIR (950–1700 nm) sensitive PMT (H12397-75, Hamamatsu). Multiphoton fluorescence, usually in visible spectral region, is reflected by the 650 nm long-pass dichroic mirror (DMLP650R, Thorlabs), passing through a band-pass filter (495-540 nm), and detected by a PMT response to visible light. For NIR-II fluorescence lifetime confocal imaging, the TCSPC module (SPC-150, Becker & Hickl GmbH) integrated in a computer will construct the NIR-II fluorescence lifetime confocal image, according to the synchronous signals from the PD and the scanning unit, as well as the electrical signals from a large-bandwidth amplifier (C5594, Hamamatsu) which amplifies the signal from PMT.

## Acknowledgments

This work was supported by National Natural Science Foundation of China (61975172, 82001874 and 61735016).

## Author contributions

J.Q. conceived the idea and provided guidance for the project. J.Z. carried out all the experiments and is the main executor. T.X.W., Y.F.L. and L.Y.C. assisted optical experiments. L.Z. performed animal operations. J.Z. and J.Q. co-wrote the paper.

