## Supplementary information for "Self-confocal NIR-II fluorescence microscopy for in vivo imaging"

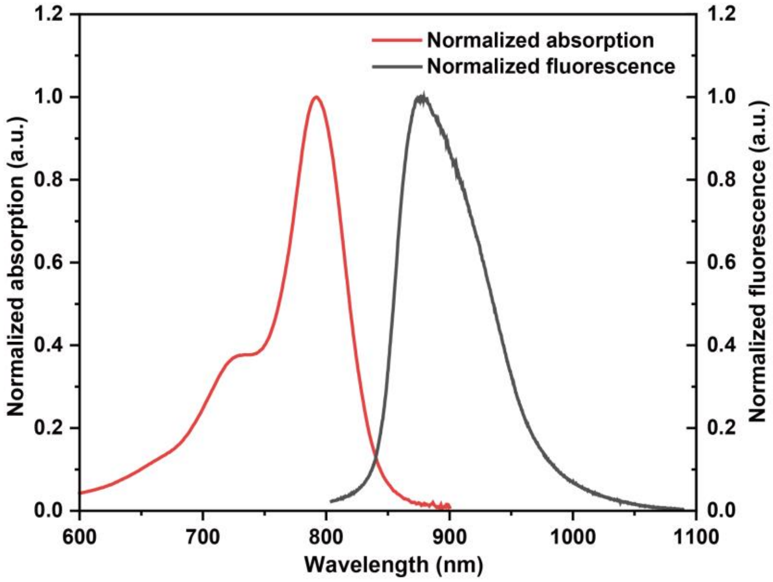


**Fig. S1** Normalized absorption and fluorescence spectra of ICG.

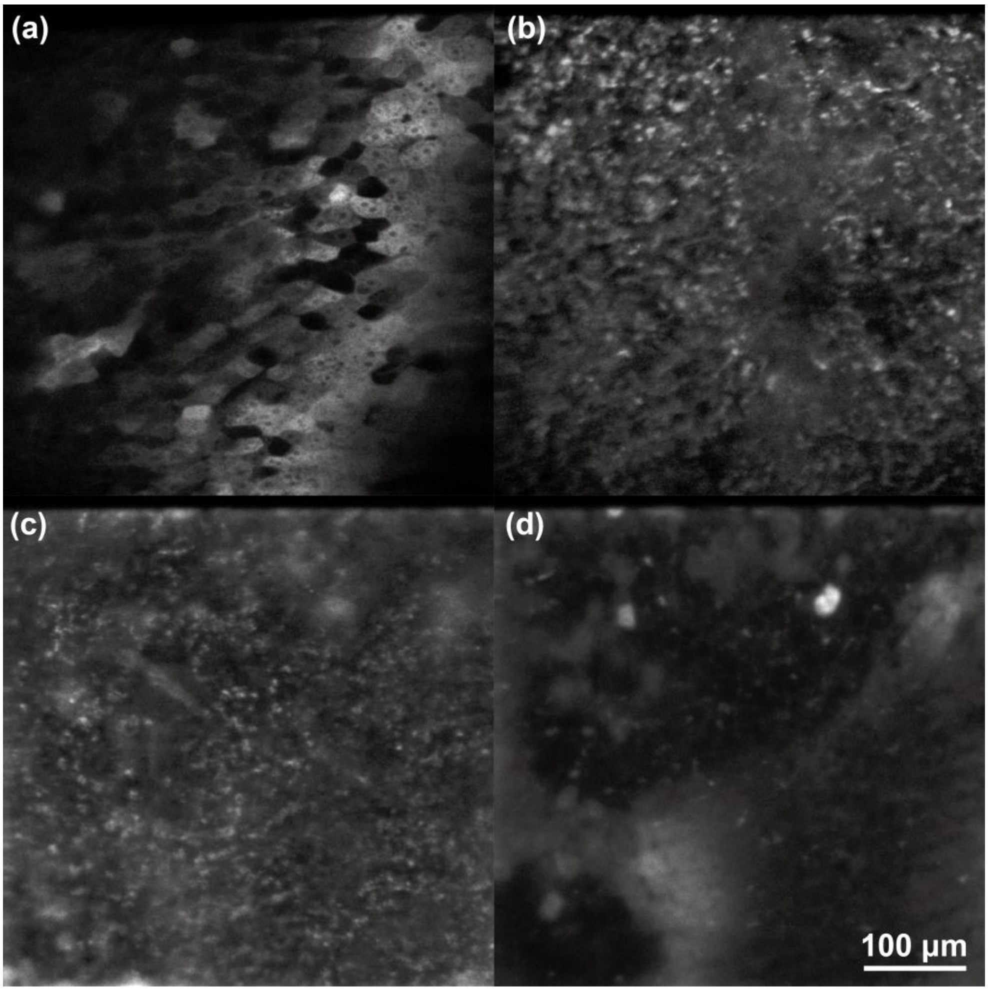


**Fig. S2** Intensity imaging of hepatic structures marked by ICG and 2TT-oC26B NPs with different NIR-II fluorescence lifetimes.
